# Age at first birth in women is genetically associated with increased risk of schizophrenia

**DOI:** 10.1101/194076

**Authors:** Guiyan Ni, Jacob Gratten, Schizophrenia Working Group of the Psychiatric Genomics Consortium, Naomi R. Wray, S. Hong Lee

## Abstract

Previous studies have shown an increased risk for a range of mental health issues in children born to both younger and older parents compared to children of average-aged parents. However, until recently, it was not clear if these increased risks are due to psychosocial factors associated with age or if parents at higher genetic risk for psychiatric disorders tend to have children at an earlier or later age. We previously used a novel design to reveal a latent mechanism of genetic association between schizophrenia and age of mothers at the birth of their first child (AFB). Here, we use independent data from the UK Biobank (N=38,892) to replicate the finding of an association between predicted genetic risk of schizophrenia and AFB in women, end to estimate the genetic correlation between schizophrenia and AFB in women stratified into younger and older groups. We find evidence for an association between predicted genetic risk of schizophrenia and AFB in women (P-value=1.12E-05), and we show genetic heterogeneity between younger and older AFB groups (P-value=3.45E-03). The genetic correlation between schizophrenia and AFB in the younger AFB group is -0.16 (SE=0.04) while that between schizophrenia and AFB in the older AFB group is 0.14 (SE=0.08). Our results suggest that early, and perhaps also late, age at first birth in women is associated with increased genetic risk for schizophrenia. These findings contribute new insights into factors contributing to the complex bio-social risk architecture underpinning the association between parental age and offspring mental health.

## INTRODUCTION

An increased risk for a range of mental health issues in children born to both younger and older parents compared to children of average-aged parents has been reported in many studies^1-8^, with particular focus on risk of schizophrenia (SCZ) in children associated with parental age^9-12^. A recent comprehensive analysis using family data extracted from the Danish Psychiatric Central Register reported a relationship between mother’s age and risk of schizophrenia in her offspring^13^. They showed that there was higher risk in children of younger and older mothers compared to those of intermediate age (25-29 years – i.e. a U-shaped relationship between maternal age and risk of SCZ in offspring), but it was unclear if this was due to psychosocial factors associated with maternal age or if mothers at higher genetic risk for SCZ tend to have children at an earlier or later age. Moreover, the very high correlation in spousal ages makes paternal and maternal contributions to this relationship difficult to disentangle. A number of possible latent mechanisms behind these epidemiological observations have been proposed^14^, including shared genetic risk factors between parents and offspring^15,16^ (Figure 1). A better understanding of factors contributing to the relationship between parental age and risk of psychiatric disorders is important for informing any future public health initiatives targeting this relationship.

**Figure 1.**
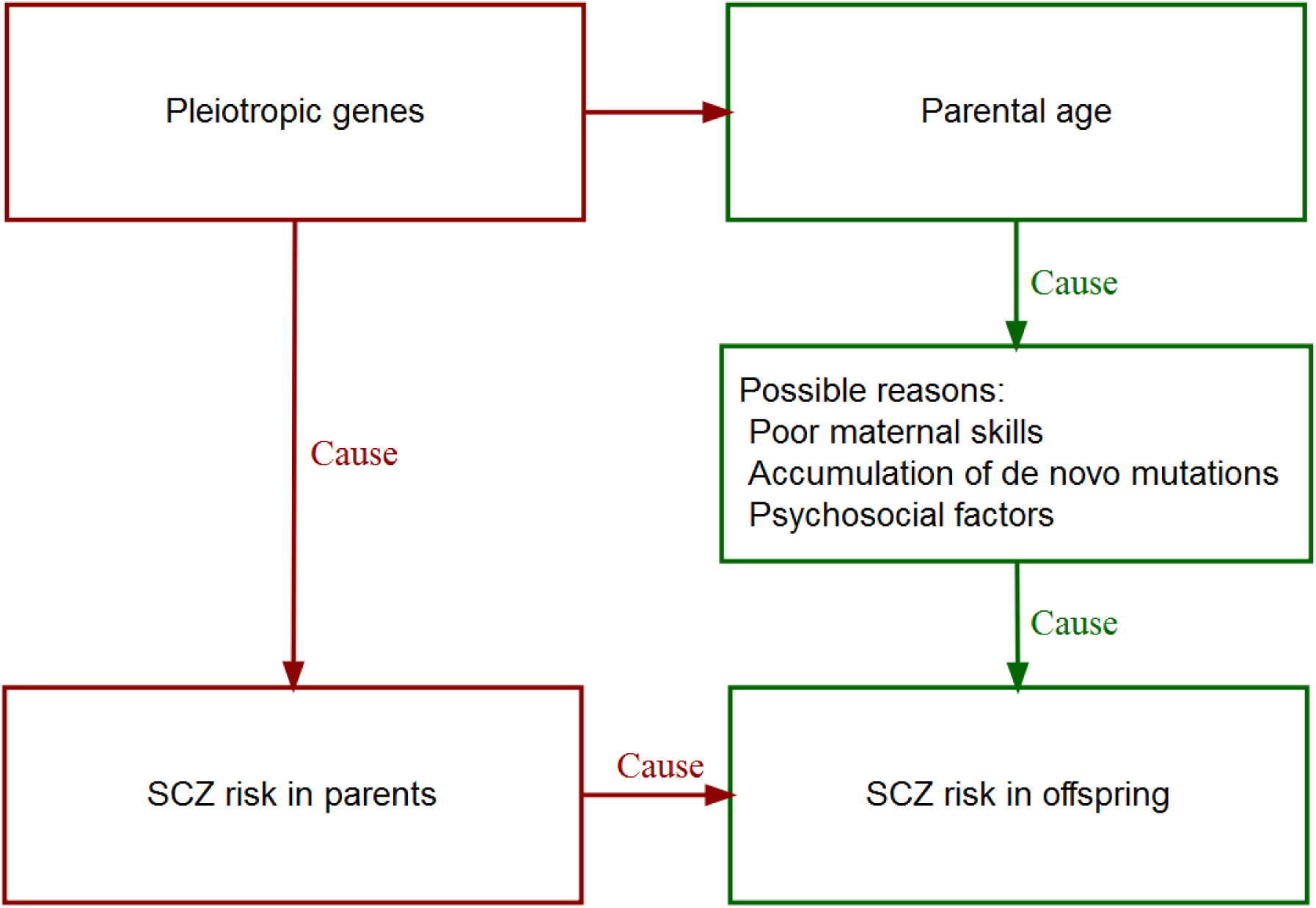
A flowchart of suggested mechanisms contributing to the relationship between the parental age and the schizophrenia risk in offspring.

We have previously reported evidence for a genetic relationship between maternal age at first birth (AFB) and risk of SCZ^17^, as illustrated in red in Figure 1. We employed a novel design that directly tests the genetic risk of schizophrenia in mothers depending on AFB. In all previous study designs the psychiatric disorder was measured in the child, and hence the relationship with AFB in the mother was confounded with characteristics of the father. Here, and in our previous study we examine the relationship between SCZ and AFB by using a genetic predictor of schizophrenia in the mother. The analyses use community samples of women enrolled in research studies that were not enriched for psychiatric disorders (< 1% for diagnosis with schizophrenia). The genetic predictor for schizophrenia can be calculated for all women in the studies as a function (such as weighted sum) of the schizophrenia risk alleles they carry, with the risk alleles identified in published genome-wide association studies (GWAS) as having increased frequencies in SCZ cases compared to controls. A woman’s genetic risk for schizophrenia is solely a function of her DNA, which she received independently of the characteristics of her partner. Hence, a benefit of this novel design is that the inferred association is not confounded by artefactual or non-genetic association(s) such as increased SCZ risk in offspring due to maternal environmental effects or by confounding with father’s age. We showed that the U-shaped relationship (between maternal age at birth and SCZ risk in offspring) observed in epidemiological studies was also observed when considering predicted genetic risk for SCZ as a function of AFB in healthy women ^13,17^.

In this study, we replicate and extend our earlier findings ^17^ using the UK Biobank data in which community samples of women have been measured for AFB. First, we confirm that SCZ polygenic risk score (PRS) for women in the UK Biobank with a record of AFB (N = 38,892) significantly predicts the U-shaped relationship found in McGrath et al.^13^, thereby replicating the results in Mehta et al.^17^. Second, we test if there is a genetic heterogeneity for AFB between younger and older AFB groups. Third, we estimate the genetic correlation between SCZ and AFB in younger and older AFB groups.

## RESULTS

### Overview

In total, 41,630 SCZ GWAS samples including 18,957 cases and 22,673 controls from 33 cohorts were used in the current study (Table S1), which were the same data used in Mehta et al.^17^. For the UK Biobank sample, 38,892 women were used in current study. The distribution of AFB, age at interview, and year of birth for the UK Biobank data after QC is shown in Figure S1. In total, 518,992 SNPs passed the quality control criteria and were in common across the SCZ and UK Biobank samples. The distribution of MAF is shown in Figure S2. Figure S3 shows that there were no closely related individuals in the UK Biobank and SCZ case-control data sets, making sure that the two data sets were independent.

We estimated SCZ polygenic risk cores (PRS) for each individual in the UK Biobank sample, using the SCZ GWAS as a reference data set (see Methods). We used both the standard genetic profile score approach^18^ (PRS-score) and the theoretically superior method of Genomic Best Linear Unbiased Prediction (PRS-GBLUP). We assessed the U-shaped relationship between AFB and SCZ PRS for the UK Biobank sample. We emphasize that in this novel design, it was not necessary to measure SCZ risk in offspring, and in our strategy potential confounding due to a correlation between paternal and maternal age was mostly removed.

Subsequently, we estimated SNP-heritability and genetic correlation between AFB and SCZ. Because of the non-linear relationship (U-shape), we divided the UK Biobank sample into two groups with younger and older AFB. We assessed if the younger and older AFB groups were genetically heterogeneous, and if there is any significant genetic correlation between SCZ and each of the younger and older AFB groups.

### Relationship between SCZ PRS and AFB

Consistent with McGrath et al.^13^ and replicating the findings in Mehta et al.^17^, a U-shaped relationship was observed between AFB and SCZ PRS-GBLUP (Figure 2). The mean SCZ PRS- GBLUP in women with early AFB (<20 years) was significantly higher than that in women with intermediate AFB (P= 2.2E-02 for AFB between 20 to < 25 years, P=1.2E-05 for AFB between 25 to < 30 years, P=2.0E-02 for AFB between 30 to < 35 years, in Table S2), but not in women with high AFB (P=4.9E-01 for AFB ≥35 years). The mean SCZ PRS-GBLUP in women with AFB between 25 to < 30 years was significantly lower than that in women with AFB between 20 to < 25 years (P=2.0E-03). Our results confirmed the findings in Mehta et al.^17^ and with stronger significance, both when PRS was calculated using GBLUP (PRS-GBLUP) and when using conventional profile scoring based on GWAS summary statistics from the SCZ GWAS data (PRS-score) (Figure S4 and Table S2). We also confirmed that the U-shaped relationship was replicated with estimated SNP effects from the full SCZ GWAS study (PRS-scorePGC) (See Figure S4 and Table S3), although these results could be biased due to possible sample overlap or the presence of relatives between the UK Biobank and the full SCZ data.

**Figure 2.**
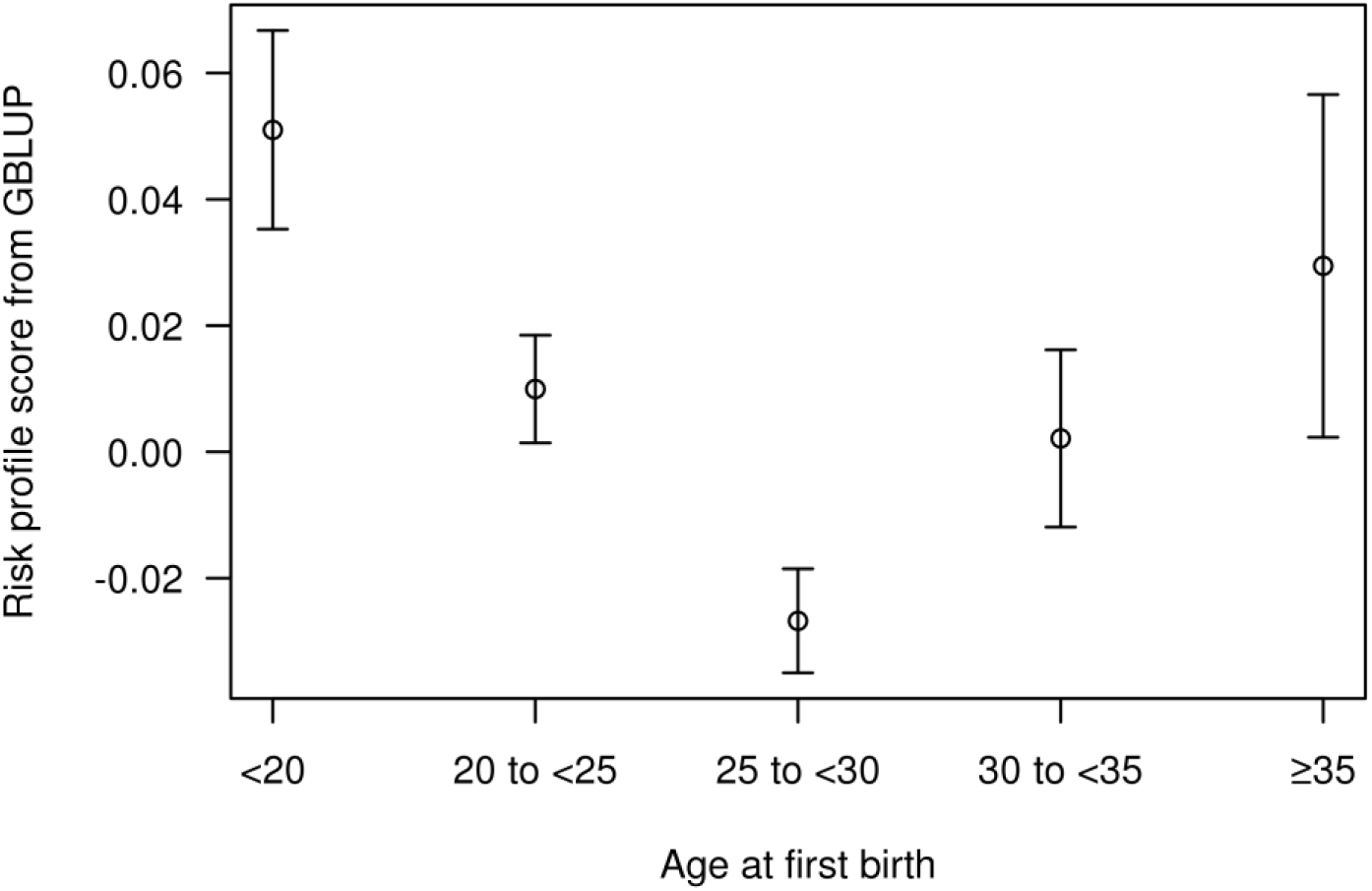
Mean and standard error of schizophrenia polygenic risk scores estimated from Genomic Best Linear Unbiased Prediction (GBLUP) in the UK Biobank sample grouped by age at first birth.

### Linear predictor

Following Mehta et al.^17^, we tested if SCZ PRS could predict the response variable that described the relationship between SCZ risk in offspring and maternal age derived in McGrath et al.^13^. Figure 3 shows that the response variable was significantly predicted by SCZ PRS for the group with the full range of AFB (P-value = 1.12E-05 for PRS-GBLUP, and P-value = 3.53E-07 for PRS-score and P-value = 3.08E-06 for PRS-scorePGC) and the subgroup with AFB younger than 26 (P-value = 4.71E-06, 6.06E-08 and 2.45E-06 for PRS-GBLUP, PRS-score and PRS- scorePGC, respectively), but not for the subgroup with AFB older than 26. The prediction with PRS-score was stronger than that with PRS-scorePGC, and both stronger than that with PRS- GBLUP.

**Figure 3.**
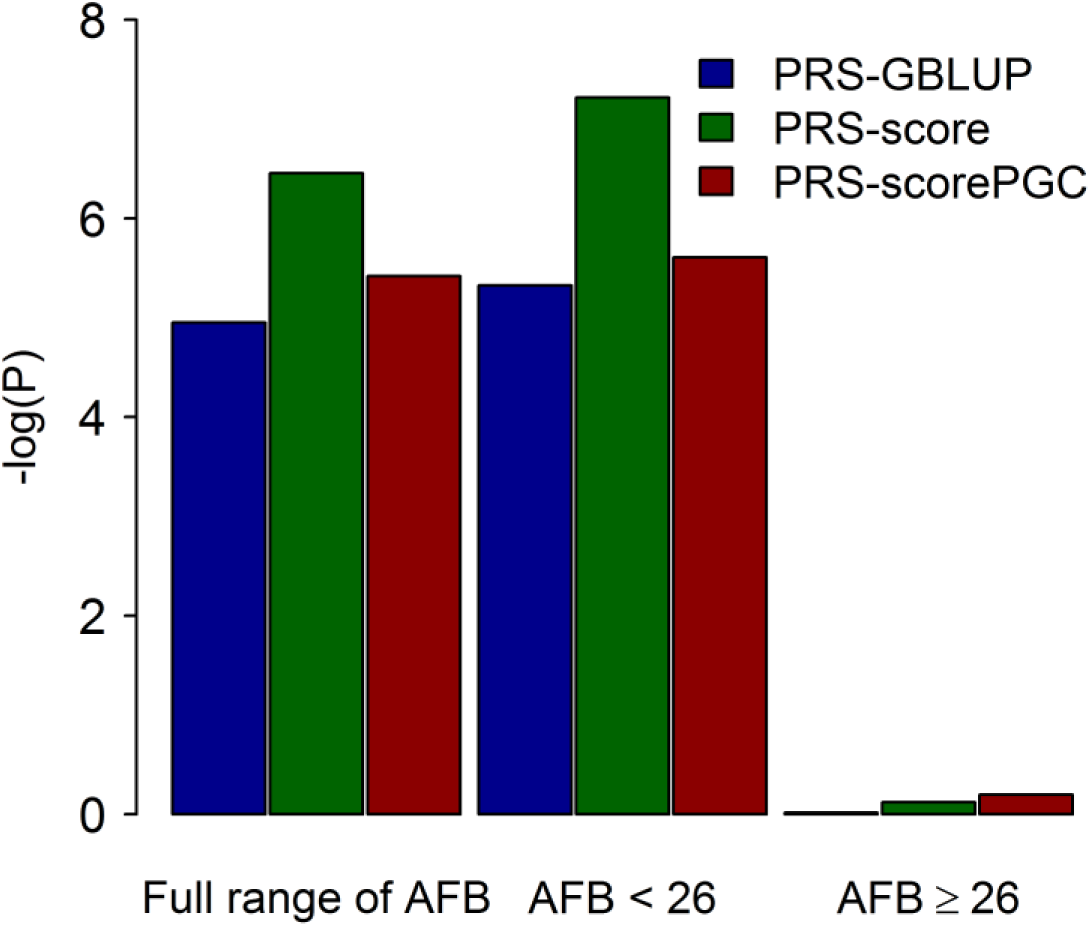
-log(P) values for the null hypothesis of R2 = 0 based on the linear prediction. Full range of AFB: All available samples with a record of age at first birth were used. AFB < 26 (≥26): Analyses were only focus on samples with AFB < 26 (≥26). PRS-GBLUP: Schizophrenia (SCZ) polygenic risk scores estimated from genomic best linear unbiased prediction were used as an explanatory variable in the model. PRS-score: SCZ polygenic risk scores estimated from genome-wide association study based on available individual genotype data were used as an explanatory variable in the model. PRS-scorePGC: SCZ polygenic risk scores estimated from summary statistics results of full PGC SCZ GWAS study were used as an explanatory variable in the model. Response variables were generated with a polynomial function derived by Mehta et al. 17, which describes the relationship between SCZ risk in offspring and maternal age (Z = 2.7214 - 0.1105X+0.0018X2, where X is age at first birth), and used in the model in which the AFB phenotypes were adjusted for age at interview, year of birth, assessment center at which participant consented, genotype batch, and the first 20 principal components.

Education level, income level, smoking and alcohol drinker status were additionally used to adjust the response variable in the linear prediction to test if those factors diminish the signals. Even with this conservative model, our results for the group with full range of AFB and the subgroup with AFB younger than 26 remained significant (Figure S5 and Table S4), albeit with reduced effect size and significance. The reduced significance might be partly explained by the reduced sample size (i.e. for full range of AFB, N= 38,892 in the base model and 31,848 in model adjusted for education and income; see Table S4). In sensitivity analyses we also restricted the sample to those recruited at age ≥45 years (N=35,451), which included the vast majority of women with a record of AFB, so that results were not biased by the exclusion of women with no reported AFB measure. We found that there was no substantive difference in our results despite the reduced sample size (Table S4 vs. Table S5).

The UK Biobank sample was divided into two subgroups born before or after 1945, a boundary of postponement of AFB based on the theory of the second democratic transition^19^. For individuals born after 1945, PRS-GBLUP significantly predicted the response variable for the group with the full range of AFB and the subgroup with AFB younger than 26, even after adjusting for socioeconomic status, and smoking and alcohol drinker status, but not for the subgroup with older AFB. For individuals born before 1945, PRS-GBLUP did not significantly predict the response variable for the group with the full range of AFB (P-value = 6.52E-02) nor the subgroup with AFB younger or older than 26 (P-value = 4.99E-02 or 4.38E-01) (Table S6). Our results agreed with the results of Mehta et al.^17^ in that SCZ PRS of women significantly predicted the response variable for the group with the full range of AFB and the subgroup with AFB younger than 26. The signals became stronger for the individuals born after year 1945.

### Genetic correlation between AFB and SCZ

Given that the AFB for women in the UK Biobank data was significantly predicted by PRS- GBLUP (P-value=1.12E-05 and R^2^=4.96E-04 in Table S4), it was of interest to estimate the genetic correlation between AFB and SCZ, which is the scaled proportion of variance that AFB and SCZ share due to genetic factors.

The SNP-heritability was 0.03 (SE=0.01), 0.10 (SE=0.01), and 0.20 (SE=0.004), for older AFB, younger AFB, and SCZ, respectively. The SNP-heritability for the older AFB group was not significantly different from 0 (Figure 4 right panel).

**Figure 4.**
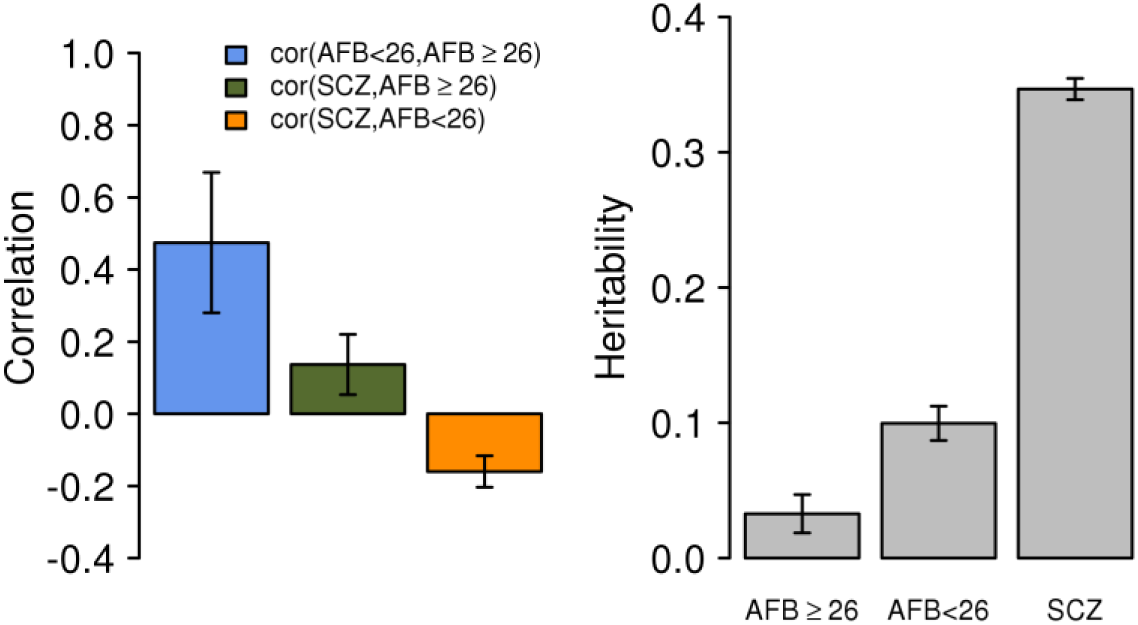
Genetic correlation (left) and heritability (right) of age at first birth (AFB) ≥ 26, AFB < 26, and schizophrenia (SCZ). Cor(AFB<26, AFB ≥26): Estimated genetic correlation between the groups with AFB < 26 and with AFB ≥ 26. Cor(SCZ, AFB≥26): Estimated genetic correlation between SCZ and AFB in the older AFB group. Cor(SCZ, AFB<26): Estimated genetic correlation between SCZ and AFB in the younger AFB group. The bars are standard errors. In the model, the AFB phenotypes were adjusted for age at interview, year of birth, assessment center at which participant consented, genotype batch and the first 20 principal components. And, the SCZ phenotypes were adjusted for sex, cohorts and the first 20 principal components. The sample size for group AFB ≥ 26 was 17,598 and for group AFB<26 was 21,294, and for group SCZ was 41,630.

The estimated genetic correlation between the younger and older AFB groups was significantly less than 1 (r^g^=0.47, SE=0.19, P-value=3.45E-03, in Table S7), indicating that younger and older AFB were genetically heterogeneous in the UK Biobank (Figure 4 left panel). We demonstrate that a truncated selection had little impact on the estimation of the genetic correlation ^20^ (Supplemental Notes 1 and 2 and Figures S6 and S7). Further, the estimated genetic correlation between SCZ and AFB in the younger AFB group was -0.16 (SE=0.04) and that between SCZ and AFB in the older AFB group was 0.14 (SE=0.08) (Figure 4 left panel).

In sensitivity analyses, education level, income level, smoking and alcohol drinker status were additionally used to adjust the phenotypes in the GREML analyses to see if those factors change the estimates. Figure S8 shows that the estimates and their significance were slightly reduced, which could be partly explained by the decrease of sample size (N=38,892 in the base model versus 31,848 in the model adjusted for education and income; see Table S7).

In addition to GREML, linkage disequilibrium score regression (LDSC)^4,21^ was applied to estimate the genetic correlation between SCZ and younger and older AFB (Table S8). As recommended by the LDSC papers^4,21^, pre-estimated LD scores from the 1000 Genome reference sample without constraining the intercept of regression were applied to the QCed GWAS data and full GWAS summary results. However, we could access individual genotype data and it was clearly known that there was no overlapping sample and no high relatedness in the QCed GWAS data for which we would be able to use LD scores estimated based on the actual genotype data as well as to constrain the intercept as zero. As reported^4,21^, it was observed from simulations (Supplemental Note 3) that if there was no overlapping sample, LDSC with constraining the intercept as zero gave the most accurate estimate with the least standard error (Figure S9). In the real data analyses, Table S8 showed that LDSC with constraining the intercept as zero gave similar estimates and standard errors, compared to those from GREML when using the QCed data. When explicitly estimating the intercept, the standard errors became large; therefore the precision of estimates might be decreased (Table S8). When using the full GWAS summary, LDSC gave similar estimates but larger standard errors, compared to GREML estimates, although a larger SCZ sample size was used for LDSC analyses (77,096 for LDSC vs. 41,630 for GREML).

## DISCUSSION

Parental age has been consistently associated with an increased risk of SCZ in offspring^9-13,15,16,22,23^, but it is well known that traditional epidemiological study designs, based on data measured for parental age and SCZ status in their offspring, have limitations with respect to disentangling genetic effects from non-genetic confounding effects such as common and residual environmental effects (Figure 1). The elevated risk of SCZ associated with parental age extends to children of both younger and older parents, compared to children of average aged parents – i.e. a U-shaped relationship^13^. The most widely accepted mechanism, in the case of delayed parenthood, is a causal relationship due to the accumulation of *de novo* mutations with age (e.g. Kong et al.^24^), although this cannot explain increased risk in offspring of younger parents. This hypothesis is biologically plausible^25-28^, but Gratten et al.^29^ have shown using theory and simulations that paternal age-related *de novo* mutations are unlikely to be the major causal factor responsible for increased risk of SCZ in offspring. Instead, they argued that increased risk of SCZ in offspring of older fathers could be due to genetic overlap between risk of SCZ and delayed parenthood. This finding is consistent with epidemiological studies showing that paternal age at first child, as opposed to paternal age at conception, accounts for the increased risk of SCZ in the children of older fathers (i.e. arguing against a direct causal role for age-related *de novo* mutations)^15,16^. Notably, this mechanism of genetic overlap between parental age and SCZ applies equally well to offspring SCZ risk associated with early parenthood.

Recently, Mehta et al.^17^ used a novel design to investigate the genetic relationship between SCZ and AFB in women that is free of many of the potential confounders present in epidemiological study designs (e.g. poor maternity skill, psychosocial factors and shared environmental factors). Specifically, they used SNP effects obtained using SCZ case-control data to estimate genetic risk of SCZ in a general community sample of women measured for AFB, finding significant evidence for pleiotropy between SCZ and AFB. In this analysis, we replicated their results in a much larger and independent community sample, the UK Biobank study. We confirmed the U-shaped relationship between AFB and SCZ PRS reported by Mehta et al.^17^ (Figures 2 and 3), and provided evidence of genetic overlap between SCZ and AFB in women. The large number of samples in the UK Biobank made it possible to also estimate genetic variance and covariance between SCZ and AFB using a linear mixed model, showing that the traits of younger and older AFB are genetically heterogeneous (Figure 4 or Table S7). The genetic correlation between SCZ and AFB in women with AFB<26 was -0.16 (SE = 0.04), which was significantly different from zero (Figure 4).

In linear prediction, results from PRS-score were similar to or more significant than those from PRS-scorePGC (Table S4). This is noteworthy because PRS-scorePGC was based on the GWAS summary statistics from the full PGC SCZ GWAS (33,640 cases, 43,456 controls), which is a larger sample than that used to generate PRS-score (individual-level genotype data on 18,957 cases, 22,673 controls). However, publicly available GWAS summary statistics such as those used for PRS-scorePGC provide incomplete information about sample overlap or pairwise relationships between data sets, either of which could introduce biases or influence statistical significance due to non-independence of samples. There is an approach or strategy for relatedness QC without accessing individual genotypes^30^, however, it is not immediately applicable to the full PGC data. We hypothesize that the superior performance of PRS-score in our analysis, despite smaller sample size than PRS-scorePGC (which is explained by restrictions on data access for individual-level genotype data), reflects the very stringent QC we applied to the data, including on relatedness.

Estimated genetic correlations and their standard errors based on GREML and LDSC were very similar (Table S8), but only when the intercept of the LDSC was constrained, which requires the strong assumption of no overlapping samples, and only when LD scores were calculated from the actual sample, which is usually not possible when using GWAS summary statistics. We also observed this phenomenon in our results based on simulated data (Supplemental Note 3 and Figure S9). These observations are in line with the statement in Bulik-Sullivan et al.^21^, that standard errors are sacrificed to achieve unbiased genetic correlation and the availability of individual-level genotype data was the most preferable scenario. Nevertheless, the estimated genetic correlations between SCZ, younger AFB and older AFB were consistent using two different approaches, i.e. GREML and LDSC (Table S8).

In summary, this study replicated previously reported evidence for significant genetic overlap between risk of SCZ and AFB in women using an independent target sample from the UK Biobank. We further showed that AFB in women is genetically heterogeneous (comparing younger to older AFB) and that there is a significant genetic correlation between SCZ and AFB. Conducting parallel analyses for AFB in men is of great interest but these data are less easily available and AFB has not been recorded for men in the UK Biobank. Our results suggest that early, and perhaps also late, age at first birth in women is associated with increased genetic risk for schizophrenia. These findings contribute new insights into factors contributing to the complex bio-social risk architecture underpinning the association between parental age and offspring mental health.

## METHODS AND MATERIALS

### Participants and quality control

#### Schizophrenia (SCZ) sample

Genome-wide association data were available from 18,987 SCZ cases and 22,673 controls from the second phase of the Psychiatric Genomics Consortium (PGC)^31^ with quality control (QC) applied as described in Mehta et al.^17^. Briefly, the SNP quality control and imputation process were performed by PGC^31^. The raw genotype data were imputed with IMPUTE2/SHAPEIT^32,33^ using the full 1000 Genomes Project dataset^34^ as the reference set. Post-imputation quality control was performed in each cohort before merging the genotype data across all cohorts, as described elsewhere^17,31^. As in Mehta et al.^17^, we used HapMap3 SNPs that were reliable in estimating (shared) genetic architecture between complex traits^35-37^. Furthermore, SNPs with call rate < 0.9, individuals with call rate < 0.9 were excluded. This less stringent QC for call rate was because SNP and individual QC had been already done by the PGC. After this QC, 688,145 SNPs and 41,630 individuals were remained and used to combine with the UK Biobank sample.

#### UK Biobank sample

In the first version of UK Biobank^38^ data set, 80,702 female (54,215 with a recode of AFB) out of 152,249 genotyped individuals were available from a community sample, in which psychiatric disorders were not enriched, with 72,355,667 imputed SNPs available.

Out of all imputed SNPs, 1,242,190 HapMap3 SNPs were identified, which were filtered through the following QC filtering criteria: SNPs with imputation INFO < 0.6^17^, minor allele frequency (MAF) < 0.01, call rate < 0.95, and Hardy-Weinberg equilibrium P-value < 10-7^17^ were excluded. In addition, ambiguous strand SNPs were excluded. After this QC, 930,841 SNPs remained and they were used to merge with the SCZ SNP list to generate a SNP-list common between the data sets. For individual level QC, only Caucasian females were used who clustered within 6 standard deviation from the mean of the EUR reference sample^39^ for the first and second genetic relationship principal components. Individuals were further excluded due to call rate < 0.95. In addition, one in a pair of individuals was randomly removed if their genomic relationship coefficient was more than 0.05^17^. An important reason for removing closely related individuals was to reduce the possibility that the similarity between those individuals could be caused by non-genetic effects (e.g. environment effects)^40^. Furthermore, UK Biobank samples were excluded if their genomic relationship with any individual in the SCZ or AFB datasets used in Mehta et al.^17^ was >0.05, in order to ensure the independence of the UK Biobank sample. The AFB sample in Mehta et al.^17^ included 12,247 genotyped women measured for AFB, who were from four cohorts: Estonia, the Netherlands, Sweden, and the United Kingdom. After QC, we used 80,522 individuals (18,957 SCZ cases, 22,673 SCZ controls, and 38,892 UK Biobank individuals) and 518,992 SNPs in the main analyses.

### Statistical analyses

#### Estimation of SCZ polygenic risk score (PRS) in UK Biobank sample

We used a GBLUP model to generate SCZ PRS for each individual in the UK Biobank sample accounting for the genetic relationship between the UK Biobank sample and the SCZ case-control sample. The GBLUP model can be written as

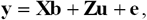

where **y** is a vector of phenotypic data (i.e. **1**s for SCZ cases, **0**s for SCZ controls and missing for UK Biobank individuals), **b** represents vectors of fixed effects including sex, cohort and 20 ancestry principal components (PCs), **u** is the vector of SCZ PRS, and **e** is the vector of residuals. **X** and **Z** are design matrices to allocate effects to phenotypic data. It is assumed that **u** is normally distributed as 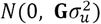, where **G** is the genomic relationship matrix constructed as described in Yang et al.^41^ and 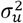 is the additive genetic variance, and **e** is normally distributed as 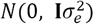, where **I** is an identify matrix and 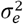 is the residual variance. GBLUP was performed using GCTA ^41^ or MTG2^42,43^ so that a subset of u for the UK Biobank sample could be inferred based on the phenotypes of the SCZ sample and the genomic relationships between the two data sets. Mean SCZ PRS in the UK Biobank individuals grouped by their AFB was estimated to assess the previously reported U-shaped relationship^13,17^. GBLUP provides more accurate PRS (PRS-GBLUP) under a polygenic genetic architecture than the more standard genetic profile score approach^18^, but for comparison we also calculated PRS by the standard method (PRS-score). To estimate SCZ risk SNP effects, an association test was conducted with PLINK 1.9^44^ using the same SCZ GWAS data used in the GBLUP analyses, with phenotypes adjusted for sex, cohort and 20 PCs. PRS-scores for individuals in the UK Biobank sample were generated by summing the count of SCZ risk alleles weighted by the SNP effects estimated from the association test. In addition to PRS-score, we used the estimated SNP effects from the full SCZ GWAS study (33,640 cases and 43,456 controls; publicly available at <u>https://www.med.unc.edu/pgc/)^31^</u> to calculate a further profile risk score in the UK Biobank sample (PRS-scorePGC), although we cannot exclude the possibility of sample overlap or relatedness between the UKB and PGC SCZ samples.

#### Linear prediction

Using 2,894,688 records from the National Danish Registry, McGrath et al.^13^ reported a U-shaped relationship between risk of SCZ in children and maternal age at birth. The resulting equation from the U-shaped relationship (z = 2.7214 - 0.1105X+0.0018X^2^) can be applied to data of age at first birth of women to generate predictors of risk of SCZ in their children^17^. We did not consider the second model in Mehta et al.^17^ that was adjusted for partner’s ages because the model was shown to be over-corrected^17^. We calculated the response variable (z) in the UK Biobank sample from the recorded age at first birth (X) and this was used as the y-variable in analyses regressing on either PRS-GBLUP, PRS-score or PRS-scorePGC and including age at interview, assessment center at which participant consented, genotype batch, year of birth and the first 20 PCs as covariates. Socioeconomic status (i.e. education and income level)^45^ or smoking and alcohol drinker status were additionally used to adjust the response variable in sensitivity analyses (Figure S1). Linear models were applied to the group with the full range of AFB records, the subgroup with AFB younger than 26 years (< 26), and the subgroup with AFB at or older than 26 (≥ 26), respectively, where the value of 26 is the mean of AFB in the UK Biobank sample. From the model, the coefficient of determination (R^2^) and P-value against the null hypothesis (i.e. SCZ PRS of women is not a predictor of AFB) were estimated.

#### Genomic residual maximum likelihood (GREML)

It is of interest to test if AFB in women is a genetically heterogeneous trait, for instance by estimating the genetic correlation between younger and older AFB groups. If the genetic correlation is significantly different from 1, it would imply that the causal variants and/or their effect sizes differ between younger and older AFB. Moreover, it would be interesting to estimate genetic correlations between SCZ case-control data and younger or older AFB groups. The UK Biobank data were divided into two groups by younger (< 26) and older AFB (≥ 26). Then, three-variate linear mixed model analysis was conducted to estimate genetic variance and covariance between SCZ case-control data, and younger and older AFB groups. The model can be written as

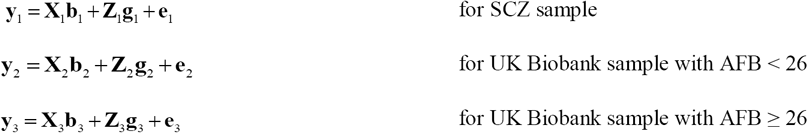

where **y** are three column vectors of phenotypic observations (one for each data set or group, i.e. SCZ case-control data set, UK Biobank AFB < 26 and UK Biobank AFB ≥ 26). For SCZ case-control data, pre-adjusted phenotypes corrected for sex, cohort and 20 PCs were used. For the UK Biobank sample, the AFB phenotypes were pre-adjusted for sex, age at interview, year of birth, assessment center, genotype batch and 20 PCs. Again, other possible confounding factors such as socioeconomic status, and smoking and alcohol drinker status were additionally controlled in sensitivity analyses to check if the estimates were substantially changed. The GREML analyses were conducted with GCTA ^41^ or MTG2^42,43^ to estimate pairwise genetic correlations among the three data sets; SCZ data, and younger AFB and older AFB.

## Acknowledgements

This research is supported by the Australian National Health and Medical Research Council (1080157, 1087889, 1103418, 1127440), and the Australian Research Council (DP160102126, FT160100229). This research has been conducted using the UK Biobank Resource. UK Biobank (<u>http://www.ukbiobank.ac.uk</u>) Research Ethics Committee (REC) approval number is 11/NW/0382. Our reference number approved by UK Biobank is 14575.

## Author contributions

S.H.L. conceived the idea and directed the study. G.N. and S.H.L. performed the analyses. N.R.W. and J.G. provided critical feedback and key elements in interpreting the results. S.H.L., N.R.W., G.N. and J.G. drafted the manuscript. All authors contributed to editing and approval of the final manuscript.

## Financial disclosures

The authors declare that they have no conflicting interests.

